# Spindle Chirp and other Sleep Oscillatory Features in Young Children with Autism

**DOI:** 10.1101/2023.06.15.545095

**Authors:** D Cumming, N Kozhemiako, AE Thurm, CA Farmer, SW Purcell, AW Buckley

## Abstract

**Objectives:** To determine whether spindle chirp and other sleep oscillatory features differ in young children with and without autism.

**Methods:** Automated processing software was used to re-assess an extant set of polysomnograms representing 121 children (91 with autism [ASD], 30 typically-developing [TD]), with an age range of 1.35-8.23 years. Spindle metrics, including chirp, and slow oscillation (SO) characteristics were compared between groups. SO and fast and slow spindle (FS, SS) interactions were also investigated. Secondary analyses were performed assessing behavioural data associations, as well as exploratory cohort comparisons to children with non-autism developmental delay (DD).

**Results:** Posterior FS and SS chirp was significantly more negative in ASD than TD. Both groups had comparable intra-spindle frequency range and variance. Frontal and central SO amplitude were decreased in ASD. In contrast to previous manual findings, no differences were detected in other spindle or SO metrics. The ASD group displayed a higher parietal coupling angle. No differences were observed in phase-frequency coupling. The DD group demonstrated lower FS chirp and higher coupling angle than TD. Parietal SS chirp was positively associated with full developmental quotient.

**Conclusions:** For the first time spindle chirp was investigated in autism and was found to be significantly more negative than in TD in this large cohort of young children. This finding strengthens previous reports of spindle and SO abnormalities in ASD. Further investigation of spindle chirp in healthy and clinical populations across development will help elucidate the significance of this difference and better understand this novel metric.

## Introduction

Sleep, the “chief nourisher of life’s feast” (1), is an essential element of physical and mental health, particularly during early development. Sleep is significantly dysregulated, however, in neurodevelopmental disorders (NDD) such as autism spectrum disorder (ASD), which manifests in behavioural sleep phenomena and changes in unique sleep oscillatory signatures (2). These are observed via electroencephalogram (EEG) and include sleep spindles and slow oscillations (SO).

Spindles, the defining feature of N2 sleep, are recurrent trains of electrical activity between 11-16 Hz. They are generated by parvalbumin-expressing (PV) GABAergic interneurons in the thalamic reticular nucleus (TRN) (3–5) and are thought to represent coordination within the thalamocortical network (6). Slow oscillations are large amplitude waves between 0.5-2 Hz of synchronized “up” and “down” states which propagate in an anteroposterior direction during N3 sleep (7). Spindles and SO, influenced by the efficiency of their coupling, are postulated to promote memory consolidation and studies have shown that entrainment of these rhythms via transcranial electrical stimulation can improve performance on working memory tasks in a variety of clinical populations (8) (9).

Importantly, both spindles and SO have shown inherent alterations in a variety of neurodevelopmental and psychiatric conditions including ASD (10) (11) (12) (13) (14) (15) (16) (17) (18) (19) (20), ADHD (21) (22) (23), schizophrenia (24) (25) (26) (27), bipolar disorder (28) and others (29).

Of increasing interest is spindle chirp, or intra-spindle change in frequency. While rarely reported, several studies have observed chirp in up to 90% of nighttime spindles (30) (31) with a predominantly negative change in frequency over the course of the spindle (32) (33) (34) (35).

As noted above for spindles and SO in neurovelopmental and psychiatric conditions, chirp has been noted to change with age (36) and to be altered in different clinical groups such as schizophrenia (27), various dementias (37) and obstructive sleep apnea (30) (38). Indeed, chirp was shown in a recent paper to be one of the strongest non-REM indicators of brain age (39).

There is a growing body of literature regarding observed sleep changes in ASD compared to age-matched healthy controls. Many of these show changes in sleep architecture (11) (12) (40), decreased spindle density and frequency (13) (14) (15) (17) and decreased SO-spindle coordination (20), though some studies failed to observe these differences (16) (41). These studies have tended to include older adolescent and adult participants; of fourteen trials reviewed the average age was approximately 12.5 years (range 1.08-27 years). As ASD involves differences in brain maturation and neuronal differentiation, these changes should be observable early in development. Indeed, spindles are well known to evolve with age (42) (39) and have been proposed as markers of brain maturation (43) (44) (45) (46).

Page et al. performed a daytime nap study of NREM sleep in children with ASD compared to typically-developing (TD) controls aged 13-30 months and found specific changes in theta, fast sigma and beta oscillations (18). Farmer et al. studied 135 children between 2-6 years old separated into three cohorts-TD, ASD, non-ASD developmental delay (DD) and reported decreased spindle density and duration in the ASD group compared to TD and DD(15). Buckley et al. assessed macro sleep architecture in an earlier, smaller version of that dataset and reported decreased total sleep time, increased N3 percentage and decreased REM percentage in ASD compared to TD.

In this present study we utilized the dataset reported in Farmer et al., enlarged following their publication (15% increase in participants), to further investigate NREM sleep neurophysiology in children with ASD, extending the list of the investigated spindle characteristics and including analyses of SO and coupling between spindles and SOs which were not performed in this sample before. Additionally, in this study, we employed automatic algorithms of spindle and SO detection utilizing all segments of NREM2 sleep in contrast to the visual detection approach limited to select periods of N2 in Farmer et. al. Finally, we explored links between the spindle metrics and behavioural scores and in an exploratory analysis tested the specificity of ASD-related alterations using the DD group.

## Methods

### Participants

Polysomnogram (PSG) and behavioural testing data of 156 children (age range 1.35-8.23 years) was obtained from the original authors (15). This dataset was obtained from individuals enrolled in an NIH study on the natural history of autism, approved by the National Institutes of Health Institutional Review Board (NCT00298246). The enrollment criteria of this sample have been previously described (15) and includes 102 individuals diagnosed with autism spectrum disorder (ASD), 24 age-matched individuals with non-autism developmental delay (DD) and 30 age-matched typically-developing (TD) individuals. To maintain a focused scope, our primary analyses included only the ASD and TD groups although we utilize the DD group in exploratory analysis to test the specificity of the alterations found in the ASD group, reported in **Supplement 1**. Autism diagnosis was based on DSM-IV-TR criteria, based on best estimate diagnosis after the administration of the Autism Diagnostic Interview–Revised (47) and the Autism Diagnostic Observation Schedule (48). TD participants were evaluated by cognitive and autism screening, including the Autism Diagnostic Observation Schedule. A medication list for included participants is available in the original publication (15); their analyses did not find interactions between medications and sleep characteristics. The current study sample includes additional ASD (n=42, n=6) and TD (n=15, n=1) relative to the Buckley et al. and Farmer et al. samples, respectively. All PSGs were rescored as described below.

Given the longitudinal nature of the parent study, this dataset included some participants with multiple PSGs (range 1-4). In these instances only the first study performed was included. Ten sleep studies from the autism group were excluded as the total sleep time (TST) was less than six hours. Additionally, one study from the ASD group was removed due to poor data quality resulting in the final sample size of 91 individuals with ASD and 30 TD participants (**Table 1**).

**Table 1.**
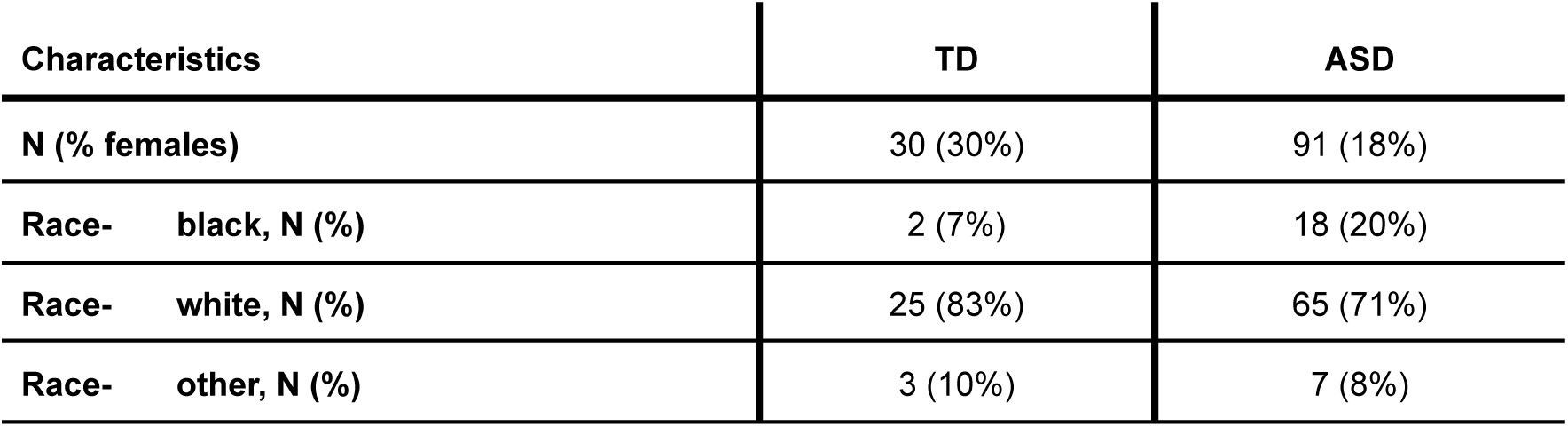
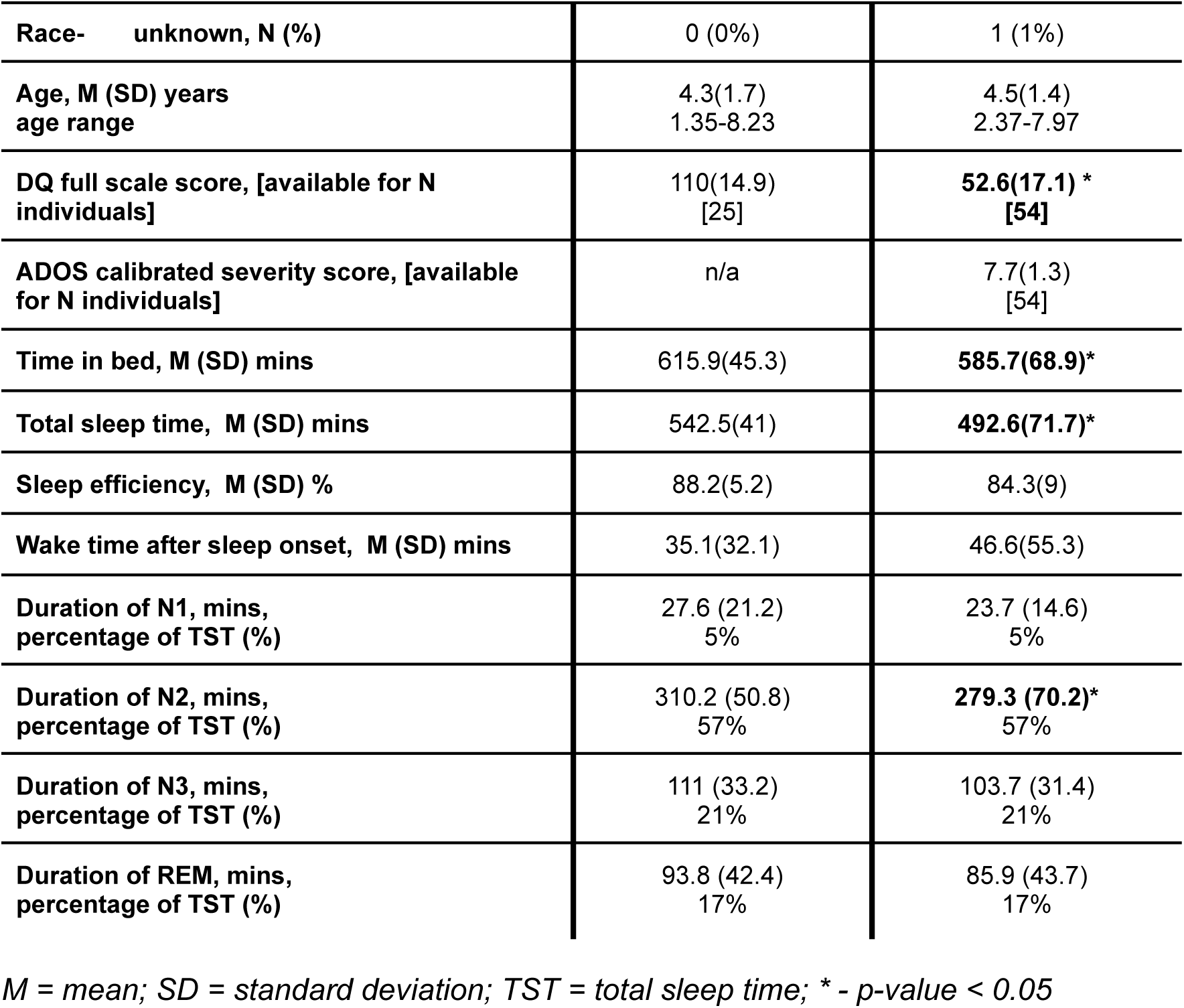
Demographics and sleep macro-architecture.

### EEG Data Acquisition

Whole-night sleep EEG data were collected from participants during an inpatient, overnight stay. As previously described (15) electrode placement followed the 10-20 system with 16 electrodes recorded (Fp1/2, F3/4, F7/8, C3/4, P3/4, T3/4, T5/6, O1/2) at 200 Hz sampling rate and referenced to contralateral mastoids.

### EEG Data Preprocessing

Sleep EEG signals were preprocessed using the open-source software Luna (https://zzz.bwh.harvard.edu/luna/). We used manual sleep stage annotations to select segments belonging to NREM2 sleep. First, we removed line noise and harmonics using a spectrum interpolation approach (49), and applied 0.5-35 Hz band-passed filter and mean-centered signals of the EEG channels. Next, we split the time series into 30-sec epochs and removed those that contained clipped or flat signals or exceeded 200 uV amplitude for 10% of an epoch length. In addition, we detected and removed outlier epochs based on Hjorth parameters (50). In the first round, outliers were defined as epochs exceeding 3 SD of the mean channel- and/or epoch-wise and in the second round, epochs with 4 SD with respect to the average across all channels and epochs were marked as outliers. Subsequently, if a particular epoch was marked as an outlier in at least 2 channels, it was removed from the analysis.

#### Sleep Architecture

Participants’ PSGs were visually scored in 30 second epochs according to AASM criteria prior to receipt of this dataset, as described in Buckley et al. 2010 and Farmer et al. 2018. Each epoch was staged as wake, REM, N1, N2 or N3. Additional metrics were assessed, including time in bed, total sleep time (TST), wake time after sleep onset, and sleep efficiency. The percentage of TST spent in each sleep stage was also calculated.

#### Spindle Detection

Following our recent reports of distinct developmental trajectories and disorder associations (51), we detected and investigated slow and fast spindles separately. They were detected using an automatic algorithm implemented in Luna specifying 11 Hz as a target frequency for slow spindles (SS) and 15 Hz for fast spindles (FS) (for a detailed description of the wavelet method of spindle detection see (51)). Spindles at each channel were characterized with respect to their average density per minute, amplitude, duration, frequency, and chirp (intra-spindle frequency change computed as a difference between mean frequencies of the first and second halves of a spindle). To investigate spindle chirp in more detail, we also estimated instantaneous frequency in an exploratory manner, using a filter-Hilbert method (bandpass filtering ±2 Hz around the mean observed spindle frequency for that individual/channel) for every sample point and subsequently averaged it for five quantiles of spindle progression (figure 2 A). Based on these estimates, we also computed the average variance and range of intra-spindle instantaneous frequency. To further visualize spindle chirp in one exemplary subject, we plot a spectrogram of averaged 2-sec epoch time-locked to spindle onset.

#### Slow Oscillation Detection

Slow oscillations were detected using an automatic approach that first 0.5-4 Hz band-pass filtered the data and detected zero-crossings. Then a series of timing and amplitude thresholds were applied to define SO. In the main analysis, we reported SO detected based on the relative amplitude threshold since it resulted in stronger coupling between SO and spindles (average magnitude of coupling between SO and SS in frontal channels 1.6 vs 1.4, p = 0.007 and between SO and FS in central channels 3.4 vs 3.2, p = 0.003). Density per minute, duration and peak-to-peak amplitude were estimated to characterize SOs at each electrode.

#### Slow Oscillation-Spindle Coupling Estimation

SO coupling with SS and FS was examined using three standard metrics: consistency of SO phase angle at spindle peak (coupling magnitude), number of spindles overlapping detected SOs (coupling overlap) and circular mean of SO phase angle at spindle peak (coupling angle). The first two metrics were additionally normalized using the mean and the standard deviation of the null distribution of each metric obtained by random shuffling (10,000 times) of the spindle peaks. In addition, we also estimated an association between SO phase angle with the instantaneous frequency of overlapping spindles as a circular-linear correlation based on eightteen 20° bins of SO phase (for more details see (27)).

### Differences from Previous Approach

As described above, our current study expands on the previous work involving this dataset (15) in that our analysis included a greater number of participants, expanded the range of spindle characteristics which were examined, evaluated SO and coupling metrics, and utilized an automated rather than manual means of spindle/SO detection. Farmer et al. manually identified and counted spindles, but were limited to five minute windows of the first four N2 periods. Using open-source software, Luna (https://zzz.bwh.harvard.edu/luna/), we were able to automatically detect spindles and SO across the entirety of N2 sleep for each PSG. Automated detection offers several distinct advantages, including increased speed of analysis and reduced error from inter- and intrascorer variability. Use of the Luna software is supported by recent work by Dickinson et al. which compared manual spindle detection to detection by Luna in children (average age 2.35 years) with and without language delay (52). In their currently unpublished work they found a strong positive correlation but poor exact agreement resulting from large mean difference (LUNA> manual) indicating that the two approaches measure related but different constructs.

### Behavioural Assessment

All participants received behavioural assessments, but only data within 90 days of the sleep study were used in the current analysis. Cognitive ability for cohort stratification was primarily assessed with the Mullen Scales of Early Learning, and the Differential Ability Scales, 2^th^ edition, or Wechsler Intelligence Scale for Children, 4^th^ edition, depending on age and ability level. In order to combine performance from both tests, developmental quotients (DQ) were calculated.

Our analyses in this present study utilized full scale DQ, which was available for 54 ASD participants and 25 TD. The DQ is interpreted in a similar fashion to an IQ, such that average performance is centered at 100. The presence and severity of behaviors associated with autism were assessed with the ADOS (Toddler module, n=36; Module 1, n = 63; Module 2, n = 52; Module 3, n = 32). To facilitate comparability across modules, we used the ordinal Calibrated Severity Score, which ranges from 1 to 10 (higher scores reflect increasing number or severity of symptoms relative to age and language level).

### Statistical Analysis

We focused our analysis only on frontal, central, parietal, and occipital channels due to the tendency of temporal and frontopolar channels to be more impacted by muscle or ocular artifacts. In addition, to reduce the number of comparisons, we averaged estimates of left and right channels resulting in estimates across four regions – frontal (mean of F3 & F4), central (mean of C3 & C4), parietal (mean of P3 & P4) and occipital (mean of O1 & O2).

A linear regression model was used to estimate the group effects (ASD vs TD) controlling for individual age, sex and race. Effect sizes of group differences were reported as differences between the group means divided by TD group standard deviation. To correct for multiple comparisons (4 regions x 19 spindles, SO and coupling metrics = 76 comparisons), we applied FDR correction with an alpha level of 0.1. To investigate potential behavioral associations between altered sleep metrics and ADOS severity score and DQ total score, we used Pearson’s correlation separately in each group and linear regression model controlling for the group, age, sex and race effects.

## Results

### Sleep Macro-Architecture

As shown in **Table 1** participants in the ASD group had significantly decreased time in bed (effect size =-0.99 SD, p=0.009) and total sleep time (effect size =-1.22 SD, p=4×10^-4^); there were no significant between-group differences in sleep efficiency or wake after sleep onset. The ASD group had significantly lower time in N2, measured in minutes (279 minutes in ASD vs 310 in TD, p=0.03), though both groups spent 57% of TST in N2. There were no between-group differences in time (in minutes) or percentage of TST spent in N1, N3 or REM.

### Spindles

There were no significant differences after multiple comparisons correction between the two study groups in slow or fast spindle density, amplitude, duration or mean frequency at any location (**Figure 1A, Sup.Table 2**). The ASD group displayed significantly more negative slow and fast spindle chirp at parietal (**Figure 1B**) and occipital regions (with effect sizes from −0.65 to −0.88 SD, all p-values < 0.01). For slow spindles, significant differences were observed in the central location (effect size = 0.55 SD, p-value=0.008) as well. In general, values of spindle chirp in both groups were lower in frontal and central leads and more positive in parietal and occipital leads.

Given the group differences in spindle chirp, which is computed here as a difference in mean frequency between the first and second half of each spindle, we further explored intra-spindle frequency change by assessing its range and variance in each group. **Figure 2A** illustrates how instantaneous frequency changes across five quantiles capturing spindle progression. For slow and fast spindles the trajectories were nonlinear in both groups and the difference in chirp was primarily driven by lower spindle frequency in the last quantiles in the ASD group compared to the typical subjects. When we compared intra-spindle frequency range and variance between the groups, they were generally comparable (except for SS intra-spindle frequency range that was higher in ASD in the central location [effect size = 0.6 SD, p-value = 0.04]) (**Figure 2B**). This suggests that a more negative chirp in the ASD group in posterior regions was not due to the increased variability in frequency over the course of a spindle.

**Figure 1.**
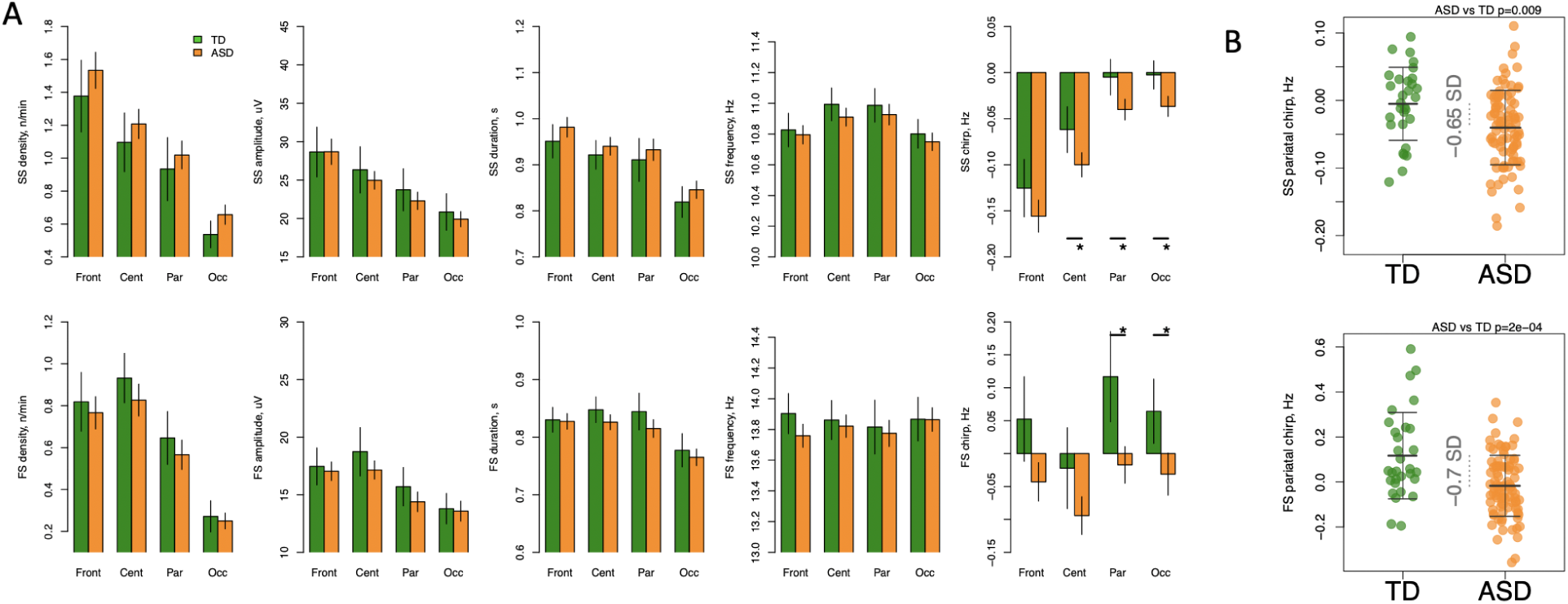
Altered spindle chirp in participants with ASD. A – spindle parameters across frontal, central, parietal, and occipital locations averaged for ASD and TD participants separately with error bars representing 95% confidence intervals. Slow and fast spindle parameters are summarized in the first and second rows respectively. B – scatterplots demonstrating group differences in spindle chirp with the largest effect sizes. FS = fast spindle; SS = slow spindle; * indicates significant differences between groups after multiple comparisons correction.

**Figure 2.**
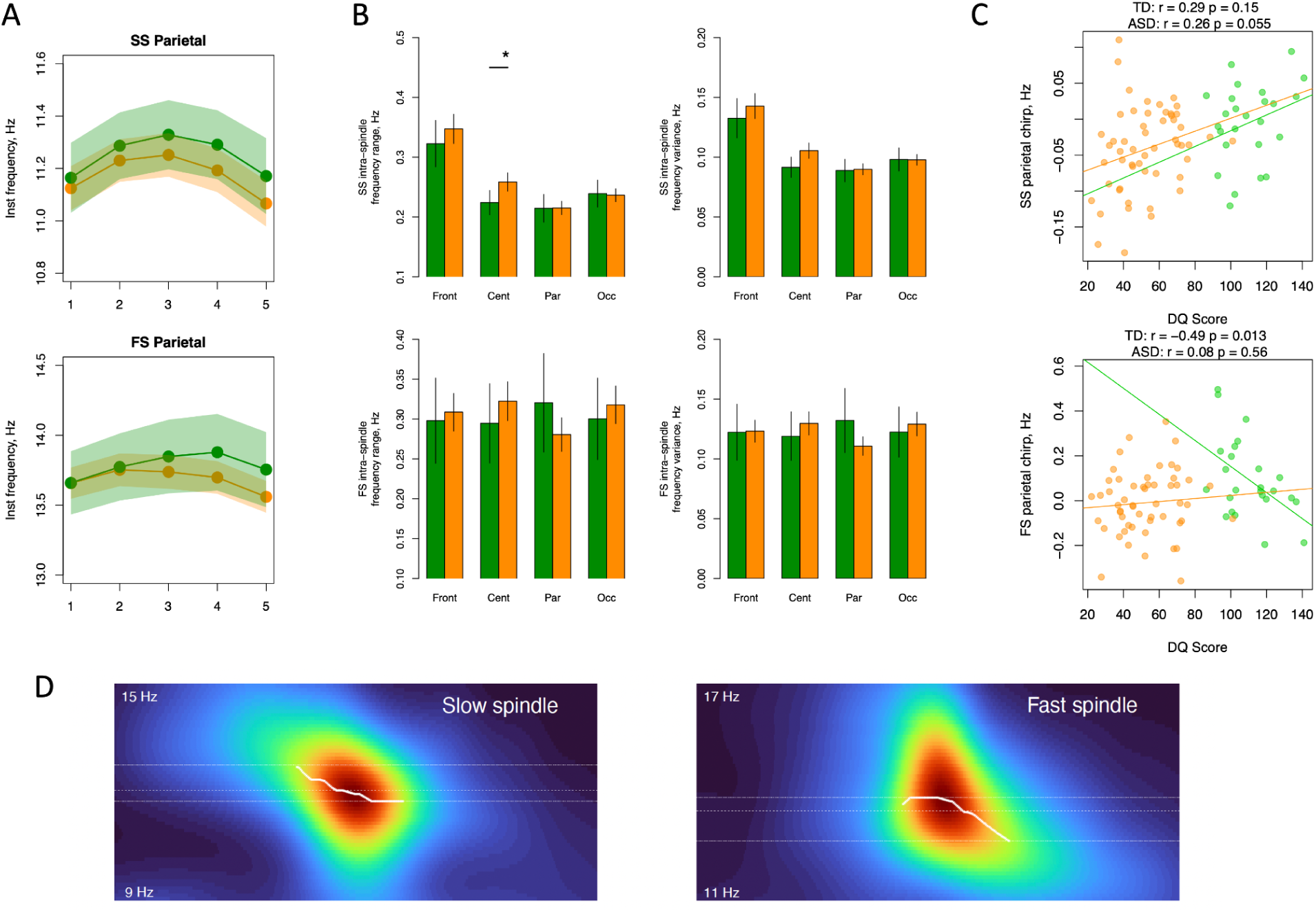
Instantaneous spindle frequency parameters. A – instantaneous frequency changes across five quantiles of spindle progression separately for ASD (in orange) and TD (in green) groups with shaded areas illustrating 95% confidence intervals. B – intra-spindle frequency range and variance over spindle progression by location. C – correlation between chirp and DQ score separately for ASD (in orange) and TD (in green) groups. D – spectrograms in sigma range time-locked to spindle onset in an illustrative subject with ASD. The white line illustrates the frequency bins with maximum power to highlight the change in frequency – chirp – over spindle progression. DQ = developmental quotient; FS = fast spindle; Inst = instantaneous (frequency); SS = slow spindle; * indicates significant differences between groups after multiple comparisons correction.

Behaviourally, we found evidence of positive association between higher values of SS chirp in parietal leads and developmental quotient score. While it did not reach the significance level in the ASD and TD groups separately (Pearson’s correlation r=0.26, p-value=0.055, r=0.29, p-value=0.15, respectively), such association was significant (p-value = 0.04) after testing it with a linear regression model controlling for the group effect, group by DQ score interaction and main covariates (sex, age and race). Concerning FS, however, the associations were different between groups. While no significant association was observed among individuals with ASD, in the TD group we found strong negative association (Pearson’s correlation r=-0.49, p-value=0.013) (**Figure 2C**). There was no significant association between SS or FS chirp and ADOS calibrated severity score in the ASD group.

We also tested if we could observe differences in chirp in the group of children with developmental delay (for full DD group description, see **Supp.Table 1**) but without ASD diagnosis and found that FS chirp values were significantly lower in the DD group compared to the TD participants in parietal and occipital locations (effect size = −0.49 and −0.45 SD, p-value = 0.008 and 0.05 respectively, **Supp. figure 1**). For SS such differences were not significant although pointed in the same direction (effect size = −0.38 and −0.28 SD). With respect to the ASD group, chirp values were more positive in the DD group but not significantly different.

### Slow Oscillations and Coupling

There were no between-group differences in SO density or duration (**Figure 3A**). The ASD group displayed significantly decreased frontal and central SO amplitude (effect sizes = −0.44 and −0.49 Sd; p = 0.004 and 0.01, respectively) with no group differences at parietal or occipital locations (**Figure 3B)**. Estimates of SO density and morphology are inherently dependent on the detection method and its settings. When an absolute SO amplitude threshold instead of the relative threshold (see methods for details) was applied, density of SO rather than their amplitude was decreased in the ASD group (max effect size = −0.55 SD, p-value = 0.002 in central region, **Supp. Fig. 2**). Thus, the results using both approaches congruently point to decreased SO activity in individuals with ASD diagnosis during N2 sleep.

**Figure 3.**
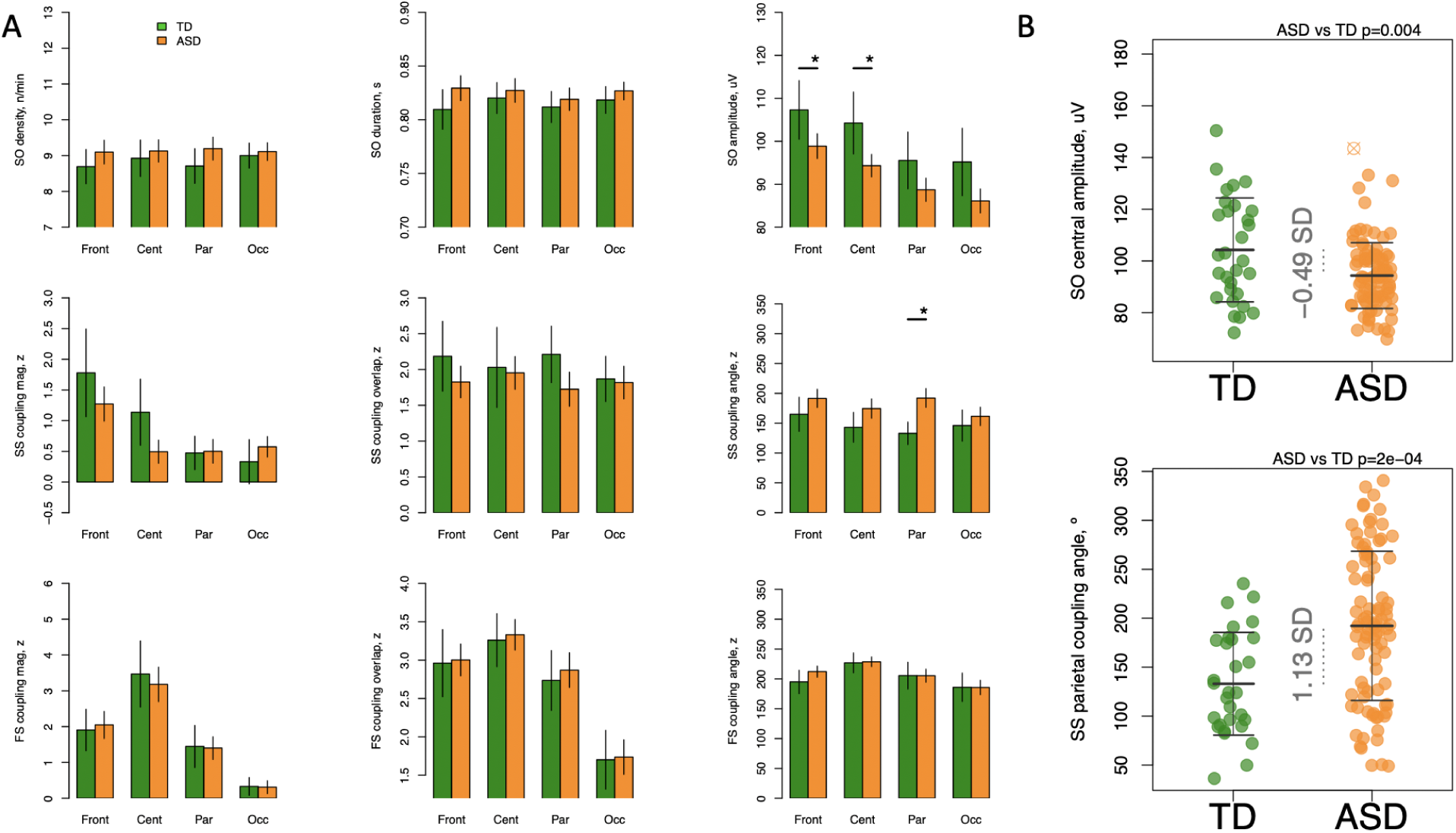
Reduced SO amplitude and altered coupling angle in ASD with preserved coupling strength. A - first row: SO density, duration, and amplitude across frontal, central, parietal, and occipital locations averaged for ASD and TD participants separately with error bars representing 95% confidence intervals; second and third rows illustrate coupling parameters between SO and slow and fast spindles, respectively. B – examples of metrics with group differences: central SO amplitude and parietal SS coupling angle. FS = fast spindle; Inst = instantaneous (frequency); SO = slow oscillation; SS = slow spindle; * indicates significant differences between groups after multiple comparisons correction.

Such reduction, however, was not linked with DQ score or ADOS calibrated severity scores in any of the groups (p > 0.05). In addition, no similar effects were observed in the DD group with respect to controls (effect sizes = 0.03 and −0.08 SD in frontal and central locations, p > 0.05, **Supp. Fig. 3**) suggesting specificity of such reduction to individuals with ASD diagnosis.

The ASD group displayed a higher parietal SO-slow spindle coupling angle (effect size = 1.13 SD, p-value = 2×10^-4^). Similar effects were also observed in frontal and central (effect size = 0.33 and 0.45 SD, p-value = 0.02 and 0.05) which were not significant after multiple comparisons correction. When an absolute SO amplitude threshold was applied, increased SO-SS coupling angle was still present with the strongest effect observed in frontal location (effect size = 0.61 SD, p-value = 0.002, **Supp. Fig. 2**). No significant association was found between DQ score or ADOS calibrated severity score and coupling angle. The DD group expressed very similar alterations with respect to the control group (effect size = 0.54 SD, p-value = 0.04 in parietal location, **Supp. Fig. 3**).

Given the differences in chirp between the groups we also tested whether we would observe differences in phase-frequency coupling – a relatively new facet of interaction between SO and spindle frequency which is measured using a circular-linear correlation. While correlation values, on average, ranged between R=0.4 and 0.55 and were slightly higher for slow than fast spindles, a comparison between the ASD group and TD group did not indicate significant group effect at any location (**Supp. figure 4**).

## Discussion

Our present study joins the body of research comparing macro and micro sleep architecture in ASD compared to age-matched TD. Similar to Kurz et al. (41) and Mylonas et al. (20) we observed a comparable distribution of sleep stages between the two groups. This is in contrast to several other studies which showed an increased SWS and decreased REM sleep ratio (40), including Buckley et al. 2010 who reported on a smaller, earlier version of this dataset. By the time of our analysis the TD group size had doubled and the ASD group increased by half, which may explain these different observations. Of note our ASD group displayed a decreased TST, similar to that in Buckley et al. 2010 and Kawai et al. 2022.

Traditionally reported spindle metrics including density, amplitude, duration and mean frequency were not significantly different between our two groups for either fast or slow spindles. While Farmer et al. (15) observed decreased spindle density and duration in a subset of this dataset, there are important study differences which may have contributed to this difference. Our present study added participants (ASD, n = 6; TD, n = 1) and measured these metrics separately for N2 fast and slow spindles across the entire night using an automated algorithm implemented in Luna software, whereas the previously reported data represented spindles as a singular entity which were visually identified in the first five minutes of the earliest four N2 periods of the night. We also utilized a different statistical approach, excluding the DD cohort from our primary analyses and included sex and race as covariates.

To our knowledge we are the first to report on spindle chirp in ASD for any age group and one of few to report on chirp in children (39). Variably referred to as chirp or intra-spindle frequency variability, it may have first been reported by Sitnikova et al. (53) in rats but has since also been observed in humans. While spindle frequency is usually reported as a mean value, studies have shown that up to 90% of spindles will display significant changes in frequency, with absolute chirp values ranging from 0.6-0.9 Hz (30) (38) (31) and it has been demonstrated to vary significantly by age (36,39). Similar to mean frequency, absolute chirp value appears to decrease after the first sleep cycle (32) (42). Studies have variably observed, or not observed, differences in chirp by scalp location or between fast and slow spindles (32) (36) (33) (34) (30) (38) (31).

In our data we observed consistently more negative chirp in the ASD group compared to TD in posterior regions for both slow and fast spindles (Figure 1). Both groups displayed a decreasing anteroposterior trend of slow spindle chirp similar to that reported by Knoblauch et al. (32) and Kozhemiako et al. (27). Fast frontal, parietal and occipital spindles in our TD group displayed a mostly positive chirp, which differs from most past studies reporting predominately negative chirp (27). However, chirp also was reported to become more negative with age over the first two decades of life (39) which may explain the more positive values in our current sample of young children.

Our exploration of instantaneous spindle frequency range and variance demonstrated similar values between ASD and TD aside from higher central slow spindle frequency range in the ASD. The similarities in variance was surprising, though it reflects similar observations by Dehghani et al. (34). It has been postulated that increased chirp in older adults represents decreased thalamocortical synchronization (36), however our findings suggest the differences in chirp within our study may be attributable to more prominent lower spindle posterior frequencies in the ASD group compared to TD.

Slow oscillations originate in the medial prefrontal cortex and project posteriorly, though it has been suggested that they have a deeper limbic source (54). Recent attention to the possible correlation between SO-spindle coupling and memory consolidation (55) (56) (9) has led to several studies assessing SO metrics in ASD. Given putative differences in numerous measures of brain maturation patterns between ASD and TD groups, we were not surprised to observe lower frontal and central SO amplitude in our ASD group. This is similar to what was reported by Mylonas et al. (20) who observed a lower SO amplitude at frontal and central locations in ASD compared to TD, though this did not reach significance in their study. These observations differ from Kurz et al. (41) who did not observe differences in SO amplitude, density or duration between ASD and TD groups. As discussed in our Results section, this difference in local amplitude was no longer significant after we applied an absolute amplitude threshold, though a significant difference in central SO density emerged TD > ASD.

Previous evaluation of SO-spindle coupling demonstrated delayed peaking in ASD with less consistent phase matching compared to TD (20). Our study showed a higher parietal SO-SS coupling angle in ASD, with a similar trend in frontal and central locations which did not survive multiple comparison correction. ASD and TD showed similar SO phase-spindle frequency coupling, indicating that spindle frequency is similarly modulated by the SO angle phase in both these groups. This supports other recent work in that, while considering a connection between SO-coupling and chirp is appealing, their effects on spindle frequency appear independent of each other (27).

Several past studies have evaluated association between sleep metrics and autism symptomatology (11) (40) (57). The sleep oscillatory features which we observed to be different between our ASD and TD groups, namely fast and slow spindle chirp, SO amplitude and SO-spindle coupling angle, did not demonstrate an association with ADOS severity score. In our analysis we did detect a positive association between parietal slow spindle chirp and a participant’s score on the full developmental quotient after controlling for sex, age and race. The original study involving this dataset collected additional behavioural data, including Vineland Adaptive Behaviour Scale and both verbal and nonverbal IQ. Having now explored and reported characteristics of advanced oscillatory signals in these cohorts, future work can include a focused analysis of the interaction between chirp and SO features and the spectrum of social functioning and cognitive ability.

Our findings from the exploratory analyses comparing the ASD and TD cohorts to a separate non-autism developmental delay (DD) group demonstrated more negative chirp than the TD group, similar to in ASD. The differences between ASD and TD SO amplitude were not observed in DD x TD or DD x ASD comparisons, and while the DD chirp values trended in a more positive direction they were not significantly different from those observed in ASD.

While spindles represent coordination between the thalamus and the cortex, recent work by Gonzalez et al. (31) looking at spindle variability between brain regions, within regions and within spindles raised interesting questions about the role of local vs global drivers of spindle dynamics. Spindle chirp, as a relatively unexplored concept, is a potentially important reflection of both local and global control and reliability of signal transmission.

As referenced above, there are numerous studies reporting observed differences in spindle and SO characteristics in ASD, however the underlying etiologies are not clear. People with autism are known to have cortical and subcortical myelin changes, including age-dependent changes in density and increased axon trajectory variability (58) (59), and myelin characteristics including overall integrity, microarchitecture and fractional anisotropy have in turn been linked to various spindle metrics (45,60–62). Within the CNS, myelin is formed by oligodendrocytes whose differentiation is driven by parvalbumin-expressing (PV) GABAergic interneurons (63). These PV interneurons are essential for coordinating synchronous cortical oscillations (64) (65) including spindles, however studies of ASD have demonstrated decreased PV interneuron density and decreased parvalbumin expression (66) (67) (59). Given the complex interplay between these factors, they may represent interesting avenues of future investigation.

In summary, our study, one of few to investigate sleep architecture in very young children with autism, observed several significant differences in the ASD cohort including more negative fast and slow spindle chirp, decreased SO amplitude and/or density depending on the employed threshold and increased SO-SS coupling angle. These differences were inconsistently associated with DQ and ADOS severity scores. While this dataset is relatively large it may have been underpowered to appreciate finer between-group differences and, while we controlled for age as a confounding factor, a larger sample size in this age group may help delineate age-specific changes between ASD and TD groups. Future studies expanding the availability of early-age PSG data in this clinical group performed alongside MRIs may help explore contributions of myelin and other changes to the differences seen in micro sleep architecture in ASD. Future investigations of spindle chirp in healthy and clinical populations of various age groups will help elucidate the significance of this metric.

## Data Availability

The data represented in this article will be made available upon reasonable request to the corresponding author.

## Supplementary Information

Participants were enrolled in the non-autism developmental delay (DD) group if their ratio IQ was at least 1.5 standard deviations below the mean, as measured by the Differential Abilities Scales (Elliot 2007) or the Mullen Scales of Early Learning (Mullen 1995) and ASD was ruled out by the Autism Diagnostic Interview–Revised and the Autism Diagnostic Observation Schedule.

---

**Suppl. Table 1.** Developmental Delay group description. (following the same inclusion criteria: all subjects had > 6 hours of TST but one subject was excluded due to persistent artifacts)

| Characteristics | DD |
| --- | --- |
| N (% females) | 23 (39%) |
| black, N (%) | 3 (13%) |
| white, N (%) | 14 (61%) |
| other, N (%) | 5 (22%) |
| unknown, N (%) | 1 (4%) |
| Age, M (SD) years<br>age range | 4.2(1.3)<br>2.35-7.59 |
| Time in bed, M (SD) mins | 601.5(74.7) |
| DQ full scale score,<br>[available for N individuals] | 59.2(14.1)<br>[12] |
| ADOS calibrated severity score,<br>[available for N individuals] | 2.4(1.3)<br>[12] |
| Total sleep time, M (SD) mins | 517.3(76.6) # |
| Sleep efficiency, M (SD) % | 86.1(8) |
| Wake time after sleep onset, M (SD) mins | 51.8(49.8) |
| Duration of N1, mins,<br>percentage of TST (%) | 24.3 (19.6)<br>5% |
| Duration of N2, mins,<br>percentage of TST (%) | 307.9 (89.2)<br>59% |
| Duration of N3, mins,<br>percentage of TST (%) | 88.4 (37.8) *<br>18% |
| Duration of R, mins,<br>percentage of TST (%) | 96.6 (54.2) 18% |

*M = mean; SD = standard deviation; TST = total sleep time, \* - p-value < 0.05 DD vs TD, #-p-value < 0.05 DD vs ASD*

**Suppl. Table 2.**
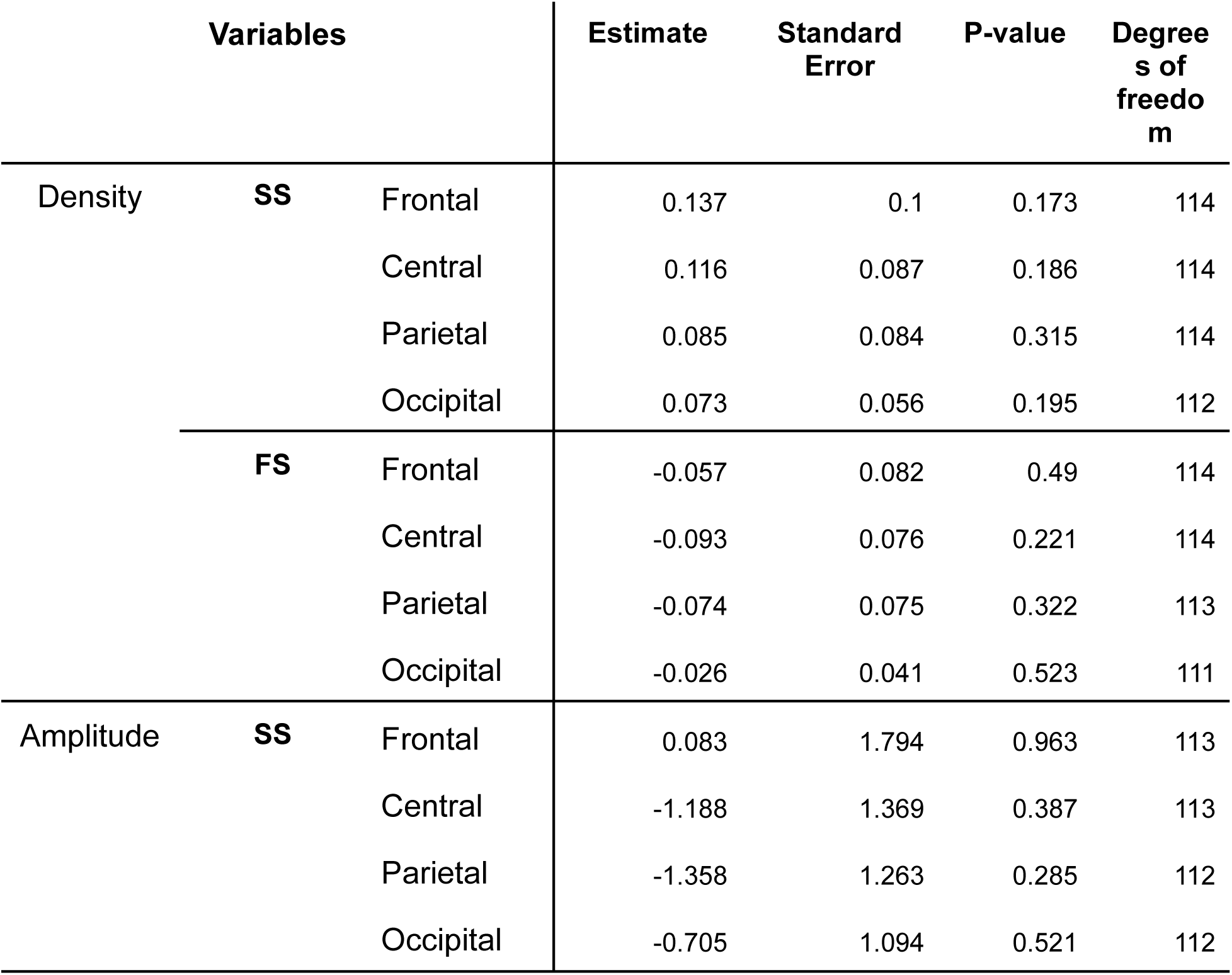
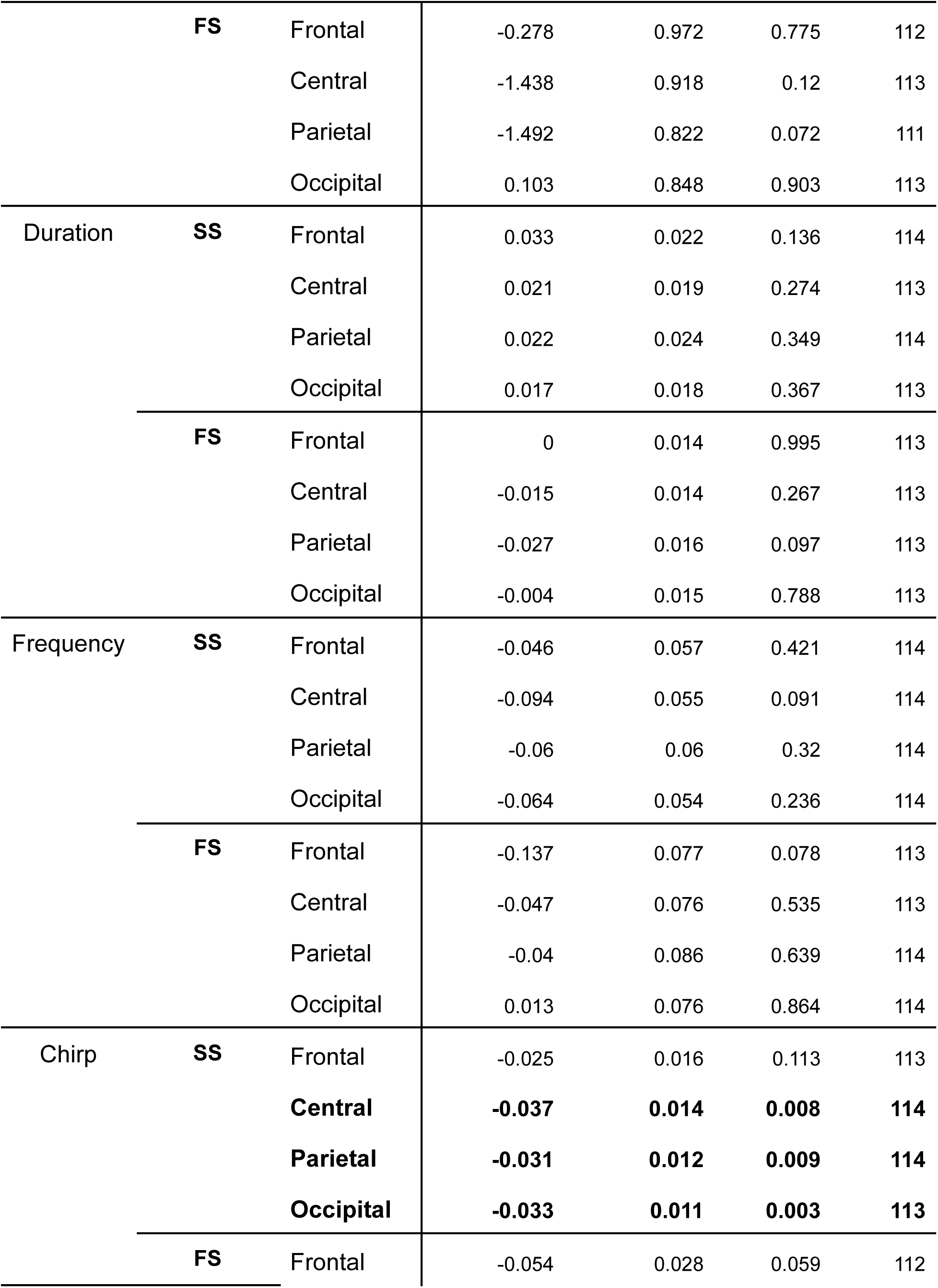
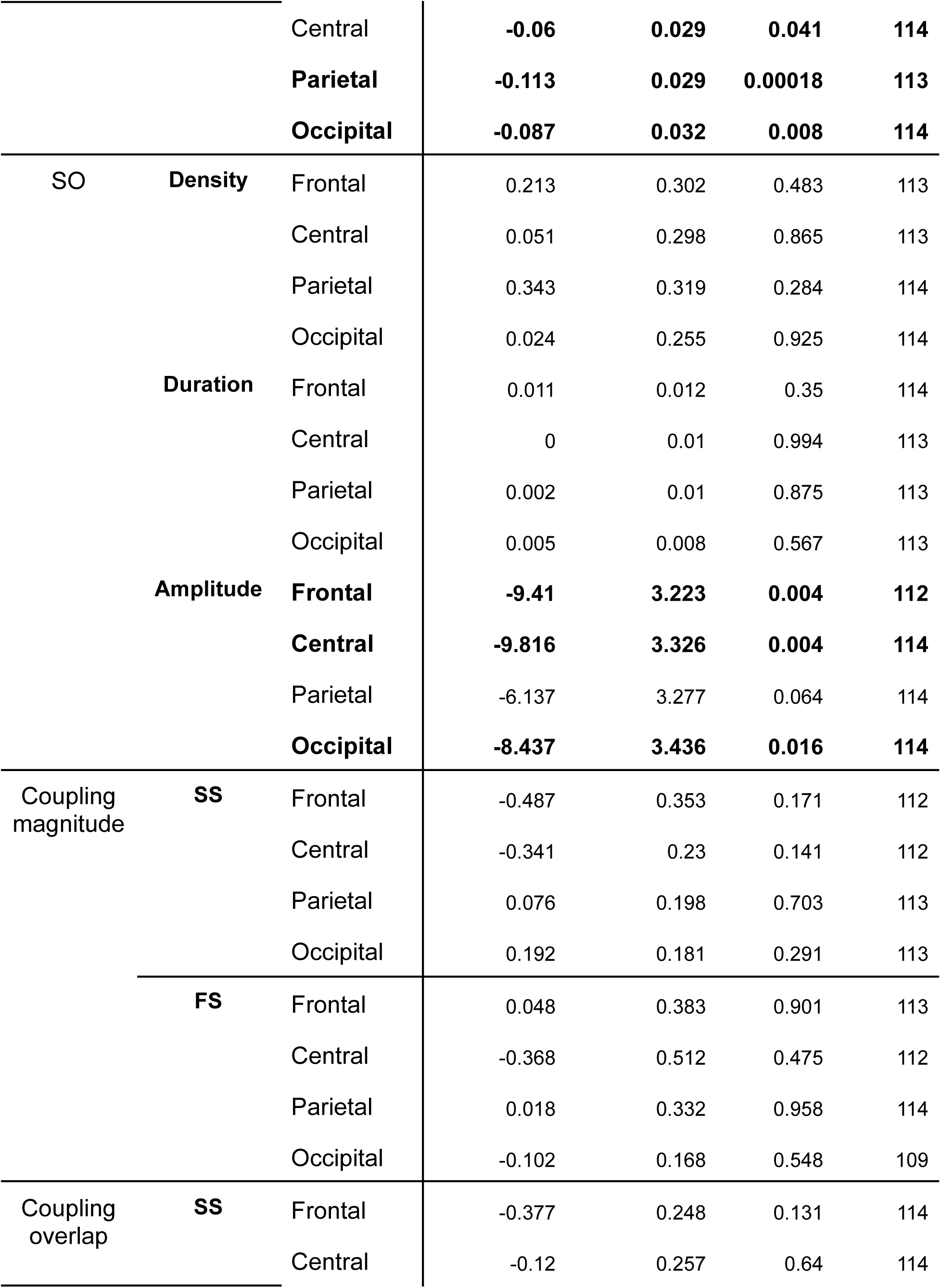
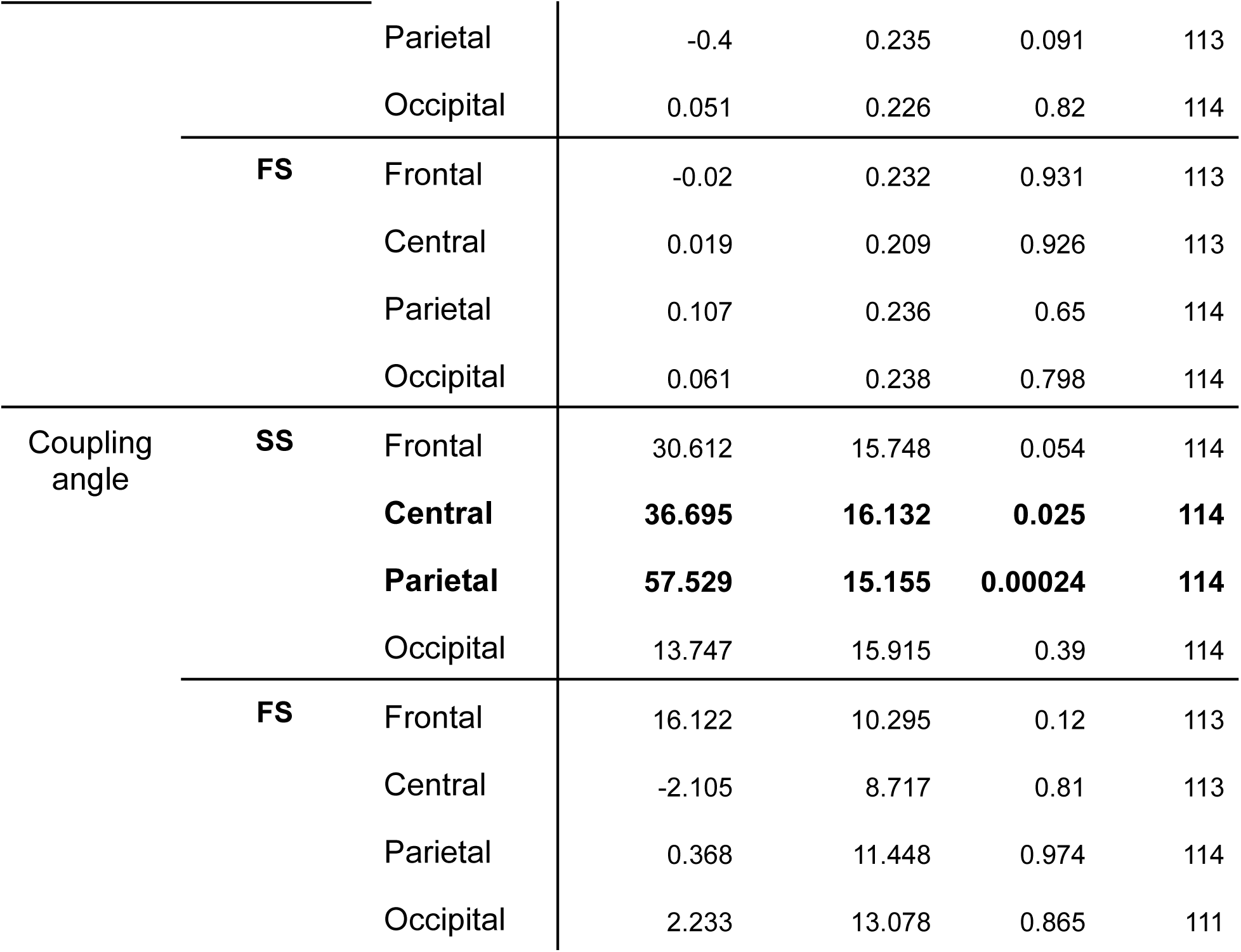
Summary statistics from liner regression model across all estimated parameters. (p-values are unadjusted for multiple comparison correction)

## Supplementary Figures

**Supplementary figure 1.**
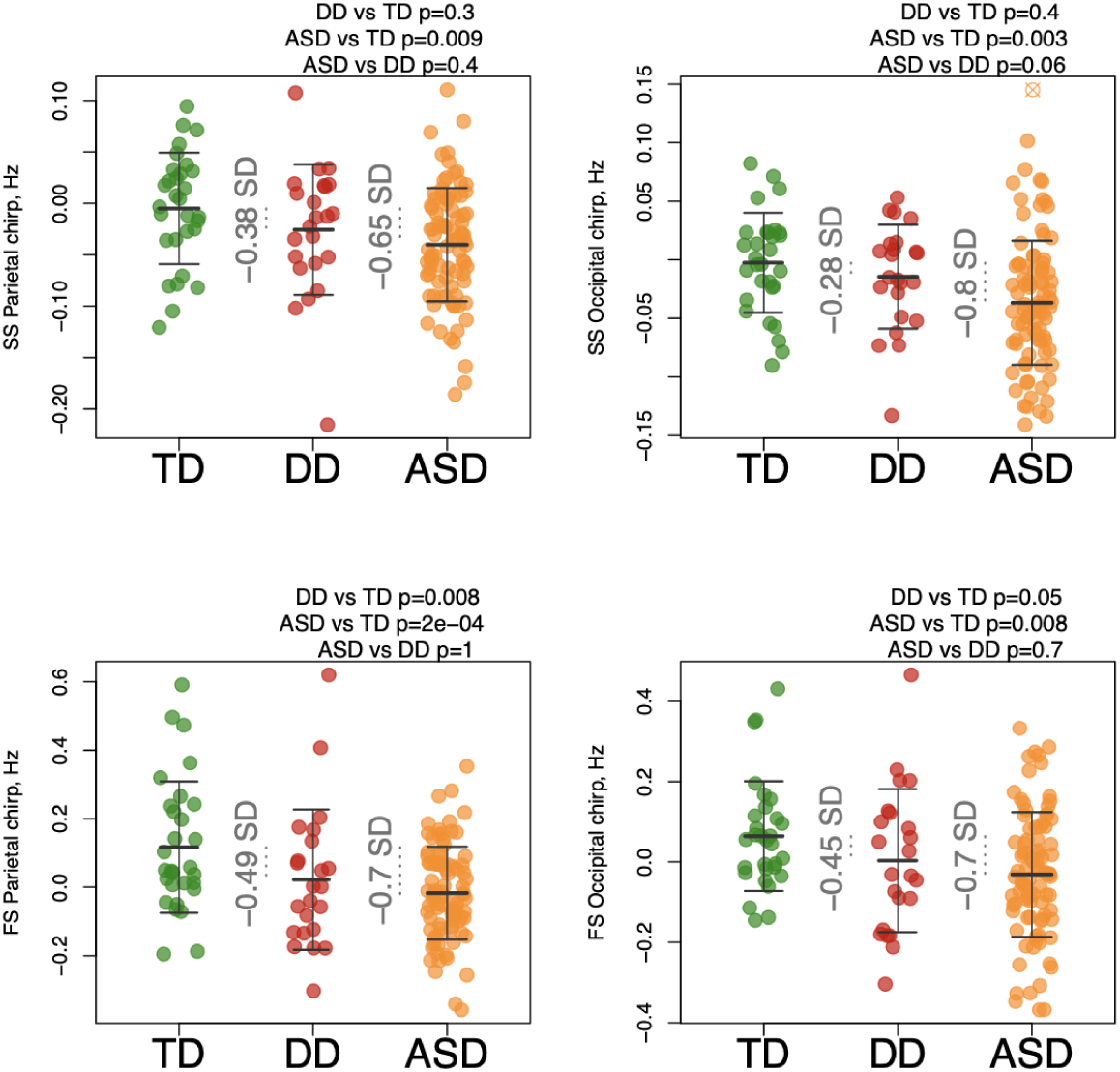
Differences in chirp in children with developmental delay. *FS = fast spindle; SS = slow spindle*

**Supplementary figure 2.**
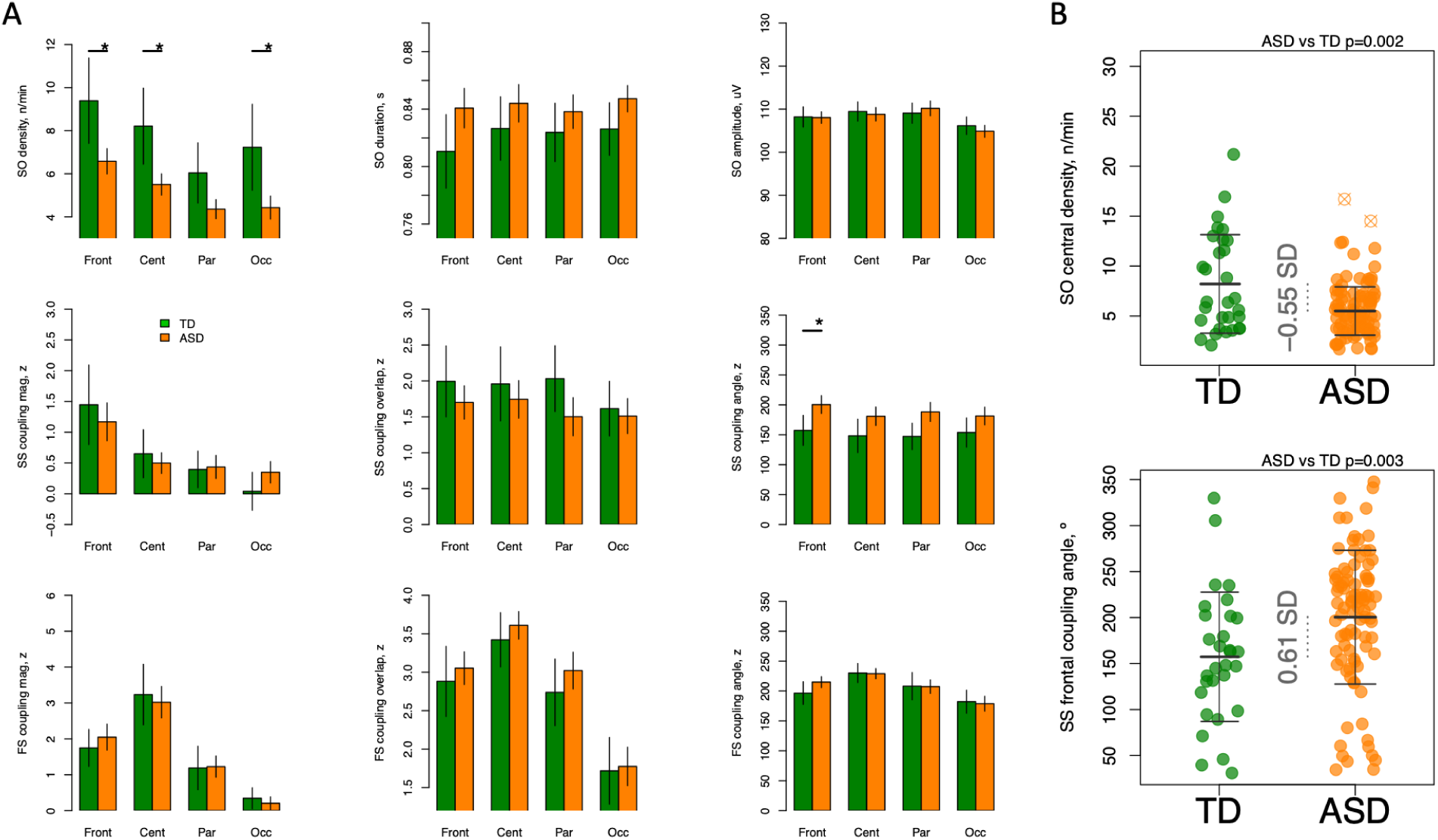
Differences in SO and coupling using absolute amplitude threshold for SO detection. *FS = fast spindle; SO = slow oscillation; SS = slow spindle; * indicates p-value < 0.05*

**Supplementary figure 3.**
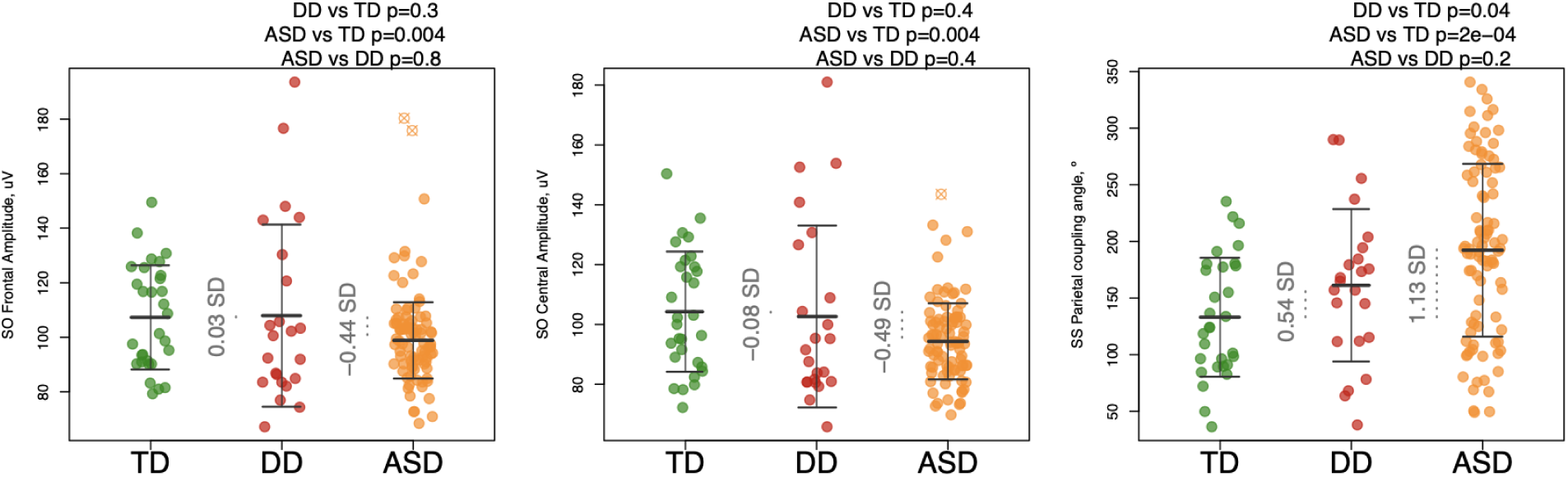
Differences in SO and coupling in children with developmental delay using relative amplitude threshold for SO detection. *SO = slow oscillation*

**Supplementary figure 4.**
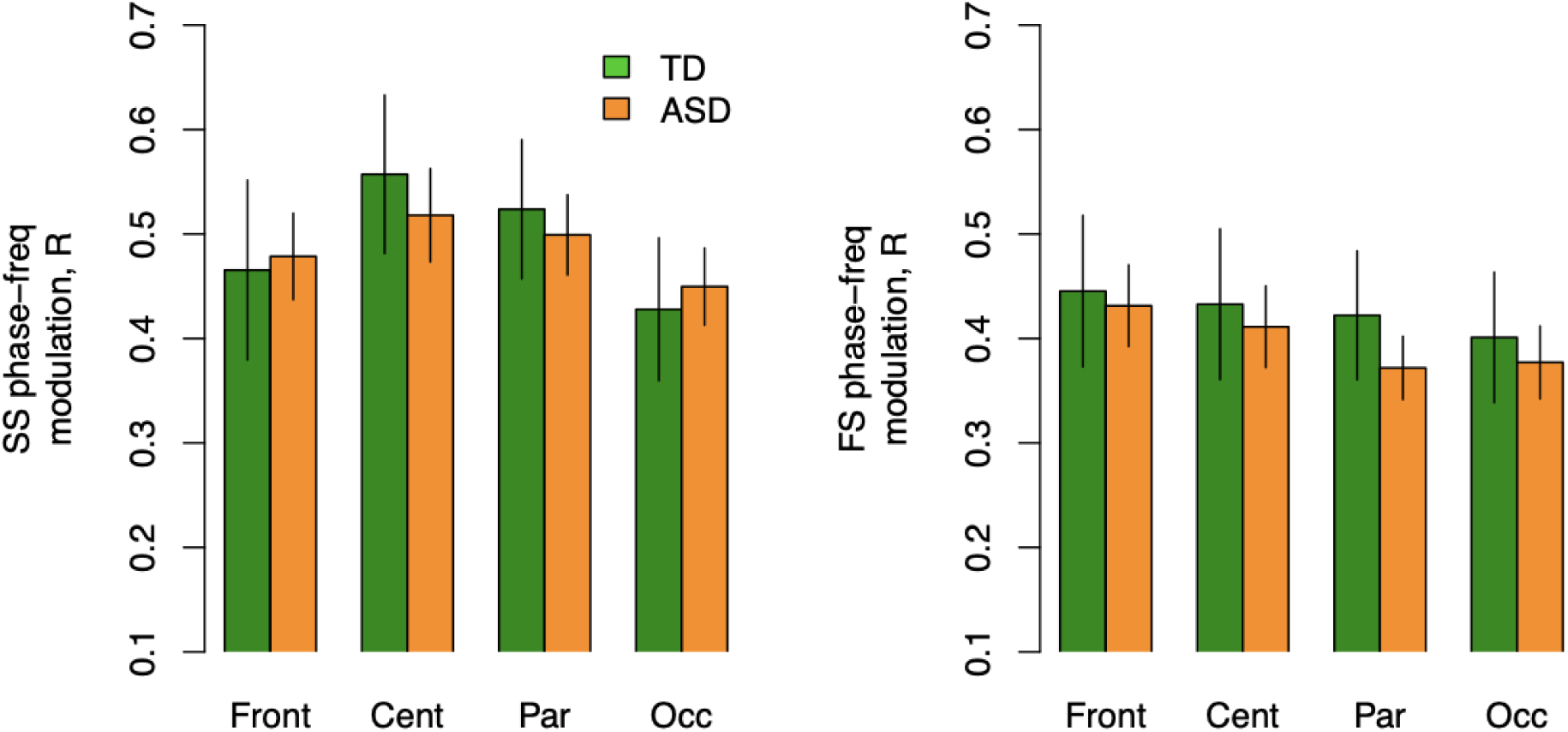
SO phase angle - spindle frequency coupling. *FS = fast spindle; SO = slow oscillation; SS = slow spindle*

52. Dickinson A, Kuhn A, Farmer C, Buckley A. Validation of an Automated Spindle Detection Method in Analyzing Neurodevelopmental Cohorts. [Unpublished manuscript].

## Suppl. References

Elliott CD. Differential Ability Scales.2nd ed. San Antonio: Psychological Corp; 2007.

Mullen EM. Mullen Scales of Early Learning. Circle Pines: AGS Publishing: 1995.

## Bibliography

1. Shakespeare W. Hamlet. East India Publishing Company; 2020.

2. Gorgoni M, Scarpelli S, Reda F, De Gennaro L. Sleep EEG oscillations in neurodevelopmental disorders without intellectual disabilities. Sleep Med Rev. 2020 Feb;49:101224.

3. Lüthi A. Sleep spindles: where they come from, what they do. Neuroscientist. 2014 Jun;20(3):243–56.

4. Clemente-Perez A, Makinson SR, Higashikubo B, Brovarney S, Cho FS, Urry A, et al. Distinct thalamic reticular cell types differentially modulate normal and pathological cortical rhythms. Cell Rep. 2017 Jun 6;19(10):2130–42.

5. Takata N. Thalamic reticular nucleus in the thalamocortical loop. Neurosci Res. 2020 Jul;156:32–40.

6. Contreras D, Destexhe A, Sejnowski TJ, Steriade M. Control of spatiotemporal coherence of a thalamic oscillation by corticothalamic feedback. Science. 1996 Nov 1;274(5288):771–4.

7. Neske GT. The Slow Oscillation in Cortical and Thalamic Networks: Mechanisms and Functions. Front Neural Circuits. 2015;9:88.

8. Barham MP, Enticott PG, Conduit R, Lum JAG. Transcranial electrical stimulation during sleep enhances declarative (but not procedural) memory consolidation: Evidence from a meta-analysis. Neurosci Biobehav Rev. 2016 Apr;63:65–77.

9. Malkani RG, Zee PC. Brain stimulation for improving sleep and memory. Sleep Med Clin. 2020 Mar;15(1):101–15.

10. Daoust A-M, Limoges E, Bolduc C, Mottron L, Godbout R. EEG spectral analysis of wakefulness and REM sleep in high functioning autistic spectrum disorders. Clin Neurophysiol. 2004 Jun;115(6):1368–73.

11. Limoges E, Mottron L, Bolduc C, Berthiaume C, Godbout R. Atypical sleep architecture and the autism phenotype. Brain. 2005 May;128(Pt 5):1049–61.

12. Buckley AW, Rodriguez AJ, Jennison K, Buckley J, Thurm A, Sato S, et al. Rapid eye movement sleep percentage in children with autism compared with children with developmental delay and typical development. Arch Pediatr Adolesc Med. 2010 Nov;164(11):1032–7.

13. Limoges É, Bolduc C, Berthiaume C, Mottron L, Godbout R. Relationship between poor sleep and daytime cognitive performance in young adults with autism. Res Dev Disabil. 2013 Apr;34(4):1322–35.

14. Tessier S, Lambert A, Chicoine M, Scherzer P, Soulières I, Godbout R. Intelligence measures and stage 2 sleep in typically-developing and autistic children. Int J Psychophysiol. 2015 Jul;97(1):58–65.

15. Farmer CA, Chilakamarri P, Thurm AE, Swedo SE, Holmes GL, Buckley AW. Spindle activity in young children with autism, developmental delay, or typical development. Neurology. 2018 Jul 10;91(2):e112–22.

16. Fletcher FE, Knowland V, Walker S, Gaskell MG, Norbury C, Henderson LM. Atypicalities in sleep and semantic consolidation in autism. Dev Sci. 2020;23(3):e12906.

17. Merikanto I, Kuula L, Makkonen T, Salmela L, Räikkönen K, Pesonen A-K. Autistic traits are associated with decreased activity of fast sleep spindles during adolescence. J Clin Sleep Med. 2019 Mar 15;15(3):401–7.

18. Page J, Lustenberger C, Frӧhlich F. Nonrapid eye movement sleep and risk for autism spectrum disorder in early development: A topographical electroencephalogram pilot study. Brain Behav. 2020 Mar;10(3):e01557.

19. Cebreros-Paniagua R, Ayala-Guerrero F, Mateos-Salgado EL, Villamar-Flores CI, Gutiérrez-Chávez CA, Jiménez-Correa U. Analysis of sleep spindles in children with Asperger’s syndrome. Sleep Sci. 2021;14(3):201–6.

20. Mylonas D, Machado S, Larson O, Patel R, Cox R, Vangel M, et al. Dyscoordination of non-rapid eye movement sleep oscillations in autism spectrum disorder. Sleep. 2022 Mar 14;45(3).

21. Gruber R, Wise MS. Sleep Spindle Characteristics in Children with Neurodevelopmental Disorders and Their Relation to Cognition. Neural Plast. 2016 Jul 11;2016:4724792.

22. Saito Y, Kaga Y, Nakagawa E, Okubo M, Kohashi K, Omori M, et al. Association of inattention with slow-spindle density in sleep EEG of children with attention deficit-hyperactivity disorder. Brain Dev. 2019 Oct;41(9):751–9.

23. Ruiz-Herrera N, Cellini N, Prehn-Kristensen A, Guillén-Riquelme A, Buela-Casal G. Characteristics of sleep spindles in school-aged children with attention-deficit/hyperactivity disorder. Res Dev Disabil. 2021 May;112:103896.

24. Ferrarelli F, Peterson MJ, Sarasso S, Riedner BA, Murphy MJ, Benca RM, et al. Thalamic dysfunction in schizophrenia suggested by whole-night deficits in slow and fast spindles. Am J Psychiatry. 2010 Nov;167(11):1339–48.

25. Manoach DS, Demanuele C, Wamsley EJ, Vangel M, Montrose DM, Miewald J, et al. Sleep spindle deficits in antipsychotic-naïve early course schizophrenia and in non-psychotic first-degree relatives. Front Hum Neurosci. 2014 Oct 7;8:762.

26. Ferrarelli F. Sleep abnormalities in schizophrenia: state of the art and next steps. Am J Psychiatry. 2021 Oct 1;178(10):903–13.

27. Kozhemiako N, Wang J, Jiang C, Wang LA, Gai G, Zou K, et al. Non-rapid eye movement sleep and wake neurophysiology in schizophrenia. eLife. 2022 May 17;11.

28. Ritter PS, Schwabedal J, Brandt M, Schrempf W, Brezan F, Krupka A, et al. Sleep spindles in bipolar disorder - a comparison to healthy control subjects. Acta Psychiatr Scand. 2018 Aug;138(2):163–72.

29. Saravanapandian V, Nadkarni D, Hsu S-H, Hussain SA, Maski K, Golshani P, et al. Abnormal sleep physiology in children with 15q11.2-13.1 duplication (Dup15q) syndrome. Mol Autism. 2021 Aug 3;12(1):54.

30. Carvalho DZ, Gerhardt GJL, Dellagustin G, de Santa-Helena EL, Lemke N, Segal AZ, et al. Loss of sleep spindle frequency deceleration in Obstructive Sleep Apnea. Clin Neurophysiol. 2014 Feb;125(2):306–12.

31. Gonzalez C, Jiang X, Gonzalez-Martinez J, Halgren E. Human Spindle Variability. J Neurosci. 2022 Jun 1;42(22):4517–37.

32. Knoblauch V, Martens WLJ, Wirz-Justice A, Cajochen C. Human sleep spindle characteristics after sleep deprivation. Clin Neurophysiol. 2003 Dec;114(12):2258–67.

33. Andrillon T, Nir Y, Staba RJ, Ferrarelli F, Cirelli C, Tononi G, et al. Sleep spindles in humans: insights from intracranial EEG and unit recordings. J Neurosci. 2011 Dec 7;31(49):17821–34.

34. Dehghani N, Cash SS, Halgren E. Topographical frequency dynamics within EEG and MEG sleep spindles. Clin Neurophysiol. 2011 Feb;122(2):229–35.

35. Schönwald SV, Carvalho DZ, Dellagustin G, de Santa-Helena EL, Gerhardt GJL. Quantifying chirp in sleep spindles. J Neurosci Methods. 2011 Apr 15;197(1):158–64.

36. Knoblauch V, Münch M, Blatter K, Martens WLJ, Schröder C, Schnitzler C, et al. Age-related changes in the circadian modulation of sleep-spindle frequency during nap sleep. Sleep. 2005 Sep;28(9):1093–101.

37. Ktonas PY, Golemati S, Xanthopoulos P, Sakkalis V, Ortigueira MD, Tsekou H, et al. Time-frequency analysis methods to quantify the time-varying microstructure of sleep EEG spindles: possibility for dementia biomarkers? J Neurosci Methods. 2009 Dec 15;185(1):133–42.

38. Souza RTF de, Gerhardt GJL, Schönwald SV, Rybarczyk-Filho JL, Lemke N. Synchronization and propagation of global sleep spindles. PLoS ONE. 2016 Mar 10;11(3):e0151369.

39. Kozhemiako N, Buckley AW, Chervin RD, Redline S, Purcell SM. Mapping Typical and Altered Neurodevelopment with Sleep Macro- and Micro-Architecture. BioRxiv. 2022 Dec 16;

40. Kawai M, Buck C, Chick CF, Anker L, Talbot L, Schneider L, et al. Sleep architecture is associated with core symptom severity in autism spectrum disorder. Sleep. 2022 Nov 17;

41. Kurz E-M, Conzelmann A, Barth GM, Renner TJ, Zinke K, Born J. How do children with autism spectrum disorder form gist memory during sleep? A study of slow oscillation-spindle coupling. Sleep. 2021 Jun 11;44(6).

42. Bódizs R, Horváth CG, Szalárdy O, Ujma PP, Simor P, Gombos F, et al. Sleep-spindle frequency: Overnight dynamics, afternoon nap effects, and possible circadian modulation. J Sleep Res. 2022 Jun;31(3):e13514.

43. Hahn M, Joechner A-K, Roell J, Schabus M, Heib DP, Gruber G, et al. Developmental changes of sleep spindles and their impact on sleep-dependent memory consolidation and general cognitive abilities: A longitudinal approach. Dev Sci. 2019 Jan;22(1):e12706.

44. Rosanova M, Ulrich D. Pattern-specific associative long-term potentiation induced by a sleep spindle-related spike train. J Neurosci. 2005 Oct 12;25(41):9398–405.

45. Piantoni G, Poil S-S, Linkenkaer-Hansen K, Verweij IM, Ramautar JR, Van Someren EJW, et al. Individual differences in white matter diffusion affect sleep oscillations. J Neurosci. 2013 Jan 2;33(1):227–33.

46. Schwichtenberg AJ, Janis A, Lindsay A, Desai H, Sahu A, Kellerman A, et al. Sleep in Children with Autism Spectrum Disorder: A Narrative Review and Systematic Update. Curr Sleep Med Rep. 2022 Nov 3;8(4):51–61.

47. Rutter M, Le Couteur A, Lord C. Autism Diagnostic Interview–Revised. Los Angeles: Western Psychological Services; 2003.

48. Lord C, Risi S, Lambrecht L, Cook EH, Leventhal BL, DiLavore PC, et al. The autism diagnostic observation schedule-generic: a standard measure of social and communication deficits associated with the spectrum of autism. J Autism Dev Disord. 2000 Jun;30(3):205–23.

49. Leske S, Dalal SS. Reducing power line noise in EEG and MEG data via spectrum interpolation. Neuroimage. 2019 Apr 1;189:763–76.

50. Hjorth B. EEG analysis based on time domain properties. Electroencephalogr Clin Neurophysiol. 1970 Sep;29(3):306–10.

51. Purcell SM, Manoach DS, Demanuele C, Cade BE, Mariani S, Cox R, et al. Characterizing sleep spindles in 11,630 individuals from the National Sleep Research Resource. Nat Commun. 2017 Jun 26;8:15930.

52. Dickinson A, Kuhn A, Farmer C, Buckley A. Validation of an Automated Spindle Detection Method in Analyzing Neurodevelopmental Cohorts.

53. Sitnikova E, Hramov AE, Koronovsky AA, van Luijtelaar G. Sleep spindles and spike-wave discharges in EEG: Their generic features, similarities and distinctions disclosed with Fourier transform and continuous wavelet analysis. J Neurosci Methods. 2009 Jun 15;180(2):304–16.

54. Morgan KK, Hathaway E, Carson M, Fernandez-Corazza M, Shusterman R, Luu P, et al. Focal limbic sources create the large slow oscillations of the EEG in human deep sleep. Sleep Med. 2021 Sep;85:291–302.

55. Marshall L, Helgadóttir H, Mölle M, Born J. Boosting slow oscillations during sleep potentiates memory. Nature. 2006 Nov 30;444(7119):610–3.

56. Ladenbauer J, Ladenbauer J, Külzow N, de Boor R, Avramova E, Grittner U, et al. Promoting sleep oscillations and their functional coupling by transcranial stimulation enhances memory consolidation in mild cognitive impairment. J Neurosci. 2017 Jul 26;37(30):7111–24.

57. Elia M, Ferri R, Musumeci SA, Del Gracco S, Bottitta M, Scuderi C, et al. Sleep in subjects with autistic disorder: a neurophysiological and psychological study. Brain Dev. 2000 Mar;22(2):88–92.

58. Ecker C, Rocha-Rego V, Johnston P, Mourao-Miranda J, Marquand A, Daly EM, et al. Investigating the predictive value of whole-brain structural MR scans in autism: a pattern classification approach. Neuroimage. 2010 Jan 1;49(1):44–56.

59. Fetit R, Hillary RF, Price DJ, Lawrie SM. The neuropathology of autism: A systematic review of post-mortem studies of autism and related disorders. Neurosci Biobehav Rev. 2021 Oct;129:35–62.

60. Mander BA, Zhu AH, Lindquist JR, Villeneuve S, Rao V, Lu B, et al. White matter structure in older adults moderates the benefit of sleep spindles on motor memory consolidation. J Neurosci. 2017 Nov 29;37(48):11675–87.

61. Gaudreault P-O, Gosselin N, Lafortune M, Deslauriers-Gauthier S, Martin N, Bouchard M, et al. The association between white matter and sleep spindles differs in young and older individuals. Sleep. 2018 Sep 1;41(9).

62. Vien C, Boré A, Boutin A, Pinsard B, Carrier J, Doyon J, et al. Thalamo-Cortical White Matter Underlies Motor Memory Consolidation via Modulation of Sleep Spindles in Young and Older Adults. Neuroscience. 2019 Mar 15;402:104–15.

63. Zonouzi M, Scafidi J, Li P, McEllin B, Edwards J, Dupree JL, et al. GABAergic regulation of cerebellar NG2 cell development is altered in perinatal white matter injury. Nat Neurosci. 2015 May;18(5):674–82.

64. Cardin JA, Carlén M, Meletis K, Knoblich U, Zhang F, Deisseroth K, et al. Driving fast-spiking cells induces gamma rhythm and controls sensory responses. Nature. 2009 Jun 4;459(7247):663–7.

65. Sohal VS, Zhang F, Yizhar O, Deisseroth K. Parvalbumin neurons and gamma rhythms enhance cortical circuit performance. Nature. 2009 Jun 4;459(7247):698–702.

66. Hashemi E, Ariza J, Rogers H, Noctor SC, Martínez-Cerdeño V. The Number of Parvalbumin-Expressing Interneurons Is Decreased in the Prefrontal Cortex in Autism. Cereb Cortex. 2017 Mar 1;27(3):1931–43.

67. Soghomonian J-J, Zhang K, Reprakash S, Blatt GJ. Decreased parvalbumin mRNA levels in cerebellar Purkinje cells in autism. Autism Res. 2017 Nov;10(11):1787–96.

